# Oxidative stress conditions result in trapping of PHF-core tau (297-391) intermediates

**DOI:** 10.1101/2020.12.07.414532

**Authors:** Mahmoud B. Maina, Youssra K. Al-Hilaly, Gunashekar Burra, Janet Rickard, Charlie Harrington, Claude Wischik, Louise C. Serpell

## Abstract

The self-assembly of tau into paired helical filaments (PHF) in neurofibrillary tangles (NFTs) is a significant event in Alzheimer’s disease (AD) pathogenesis. Oxidative stress, which accompanies AD, induces multiple post-translational modifications in proteins including the formation of dityrosine (DiY) cross-links, previously observed for Aβ. Here, metal catalysed- and ultra-violet oxidation were used to study the influence of DiY cross-linking on the self-assembly of the PHF-core tau fragment. We report that DiY cross-linking facilitates tau assembly into tau oligomers and prevents their elongation into filaments. The DiY cross-linked tau oligomers were not toxic to cells. Our findings suggest that DiY cross-linking of pre-assembled tau, promotes the formation of soluble tau oligomers that show no acute toxicity.

## Introduction

Alzheimer’s disease (AD) is the most common form of dementia, characterised by the deposition of extracellular amyloid-beta (Aβ) plaques and intracellular neurofibrillary tangles (NFTs). NFTs are comprised of paired helical filaments (PHFs) and straight filaments (SFs) composed of tau (1, 2). The burden of NFTs shows correlation with the extent of pathology in AD (3), and NFT accumulation provides a reliable staging of the disease process (4). The formation of filaments involves the self-assembly of tau monomers but the precise mechanisms involved are not fully understood. Some types of oligomers are likely to convert into to PHFs and SFs (5). The different forms of abnormal tau aggregates have been associated with specific dysfunctions associated with memory deficits: cellular toxicity (5); synaptic dysfunction; mitochondrial disturbance, and the spread of tau pathology across the brain in mouse models (6–9). Tau oligomers have also been identified in the early stages of AD (10).

The self-assembly of tau is influenced by a myriad of post-translational modifications (11). For example, while tau phosphorylation and truncation are thought to contribute to its assembly (11), the nitration of tyrosine residues has been shown to inhibit the assembly of arachidonic acid-induced tau into filaments (12, 13). Multiple studies have shown that oxidative stress generates tau post-translational modifications including phosphorylation (14, 15), acetylation (16, 17), nitration, nitrotyrosine and dityrosine (DiY) formation (12). Oxidative stress is one of the earliest sources of damage identified in the brain in AD (18) and key markers of oxidative stress, such as nitrotyrosine and DiY levels, are increased in the AD brain (19). DiY cross-linking results in a stable, irreversible modification (20). Aβ and α-synuclein (associated with Parkinson’s disease) have both been shown to form DiY cross-links in *vitro* (21–24). Modification tyrosine residues by oxidative stress has been previously demonstrated to influence tau aggregation (12, 25). For example, it has been shown that oxidative stress induced by peroxynitrite (ONOO^−^), results in the oligomerisation of full-length human tau stabilised via DiY cross-linking of arachidonic acid-induced tau aggregation (12). Although this indicates that DiY forms in tau, it is not clear whether DiY formation promotes tau self-assembly into filaments or stabilises tau oligomers *in vivo*. Tau is usually induced to form filaments with the help of heparin or other polyanionic molecule (26–30). A truncated tau fragment, corresponding to residues Ile297–Glu391, was first isolated from the proteolytically stable core of paired-helical filaments (PHFs) (2) and more recently was found to overlap with the region identified in the PHF and SF core by CryoEM (31, 32). This fragment is known as dGAE and was recently shown to self-assemble to form form filaments in vitro without the addition of exogenous additives such as heparin, RNA or arachidonic (33, 34).

The longest isoform of tau with 441 amino acid residues contains five tyrosine residues, located at positions 18, 29, 197, 310, and 394. The dGAE fragment contains a key region thought to be important for assembly competence ^306^VQIVYK^311^ (35, 36), which contains Y310. Therefore, the dGAE fragment serves as an excellent *in vitro* model to investigate the influence of oxidative conditions on DiY formation within tau and to characterise its propensity to assemble in the absence of exogeneous additives. Using metal-catalysed oxidation (MCO) and ultra-violet (UV) - induced photo-oxidation to induce DiY cross-linking, we show that the oxidation of soluble dGAE facilitates assembly into Thioflavin S (ThS)-negative Tau oligomers that lack β-sheet structure and which do not elongate into fibrils. Moreover, our results revealed that, at a higher (10:1) Cu^2+^: dGAE ratio, Cu^2+^ alone (without H_2_O_2_) facilitates the formation of these DiY cross-linked Tau oligomers, prolongs the oligomer half-life and inhibits further assembly into fibrils. We report that, unlike other tau oligomers previously reported (37–40), the DiY cross-linked tau oligomers are not acutely toxic to differentiated neuroblastoma cells in the timeframe studied. Our findings suggest that DiY cross-linking on soluble dGAE promotes the formation of non-toxic soluble tau oligomers incapable of further elongation into fibrils.

## Materials and Methods

### Preparation of dGAE tau fragment

The preparation of recombinant dGAE (tau 297-391) has been described previously (33). Briefly, dGAE was expressed in bacteria and, following heat treatment, purified by P11 phosphocellulose chromatography. 2-(N-morpholino)ethanesulfonic acid (MES), pH 6.25, was used instead of piperazine-N,N′-bis(ethanesulfonic acid) (PIPES) was used in some cases. The protein fractions were eluted with 50 mM PIPES (pH 6.8) or 50 mM MES (pH 6.25), both supplemented with 1 mM EGTA, 5 mM ethylenediaminetetraacetic acid (EDTA), 0.2 mM MgCl_2_ and 5 mM 2-mercaptoethanol containing 0.1–1 M KCl. The peak of protein elution was identified by protein assay (at 0.3–0.5 M KCl) and dialyzed against 80 mM PIPES buffer (pH 6.8), 1 mM EGTA, 5 mM EDTA, 0.2 mM MgCl_2_, 5 mM 2-mercaptoethanol, or PB (10 mM; pH 7.4). The dGAE protein concentration. was measured using Advanced Protein Assay Reagent (Cytoskeleton, Inc.) with bovine serum albumin as a standard. The protein was diluted with 10 mM phosphate buffer (pH 7.4) and for all the experiments, the dGAE was used at a concentration of 100 μM (Table 1).

**Table 1:**
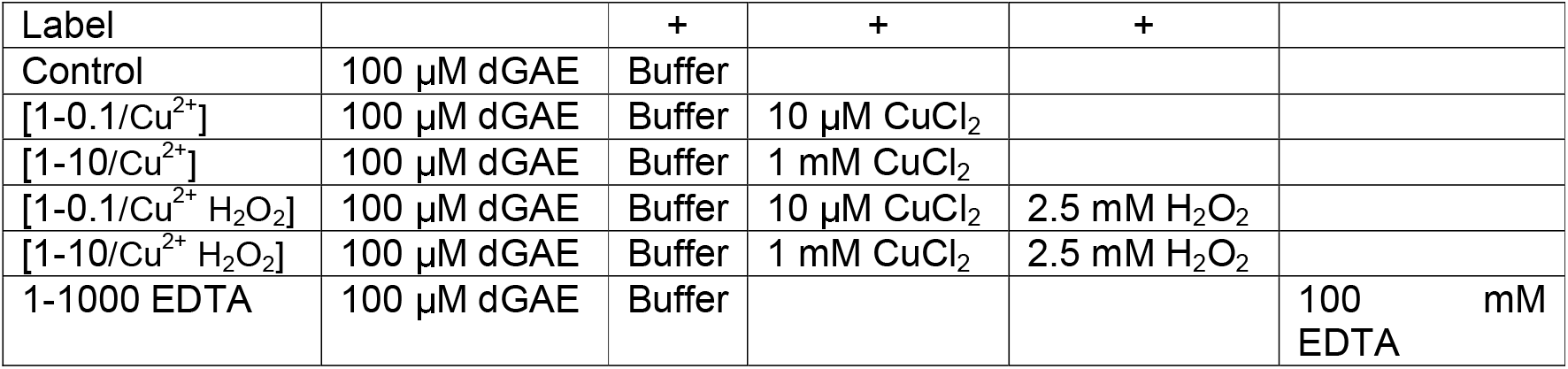
dGAE conditions with and without oxidation

| Label |  | + | + | + |  |
| --- | --- | --- | --- | --- | --- |
| Control | 100 μM dGAE | Buffer | | | |
| [1-0.1/Cu<sup>2+</sup>] | 100 μM dGAE | Buffer | 10 μM CuCl<sub>2</sub> | | |
| [1-10/Cu<sup>2+</sup>] | 100 μM dGAE | Buffer | 1 mM CuCl<sub>2</sub> | | |
| [1-0.1/Cu<sup>2+</sup> H<sub>2</sub>O<sub>2</sub>] | 100 μM dGAE | Buffer | 10 μM CuCl<sub>2</sub> | 2.5 mM H<sub>2</sub>O<sub>2</sub> | |
| [1-10/Cu<sup>2+</sup> H<sub>2</sub>O<sub>2</sub>] | 100 μM dGAE | Buffer | 1 mM CuCl<sub>2</sub> | 2.5 mM H<sub>2</sub>O<sub>2</sub> | |
| 1-1000 EDTA | 100 μM dGAE | Buffer | | | 100 mM EDTA |

### Metal-catalysed oxidation of dGAE

dGAE in 10 mM phosphate buffer, pH7.4 was prepared according to Table 1.

The control incubation with EDTA was to chelate trace metals in the reaction. All the peptides were either incubated at 37°C, or at 37°C with agitation at 350 RPM (Thermomixer C, Epppendorf, Germany). At each time point measured, the oxidation reaction was stopped using EDTA at a final concentration of 2 mM. A minimum of three independent experiments were repeated to ensure the reproducibility of the findings.

### Photo-oxidation of dGAE

dGAE at 100 μM in 10 mM phosphate buffer was incubated i) without UV-C in the dark, and ii) in the presence of UV-C for 2h using a G6T5 Germicidal 9′ 6W T5 UVC lamp set to 8 J/m^2^/sec (General Lamps Ltd). A minimum of three independent experiments were repeated to ensure the reproducibility of the findings.

### Fluorescence spectroscopy

To monitor the formation of DiY, fluorescence intensity at 400-420 nm was measured using a fluorescence spectrophotometer (Varian Ltd., Oxford, UK), using a 1-cm path length quartz cuvette (Starna, Essex, UK). Data was collected using a fluorescent excitation wavelength of 320 and emission collected between 340 – 600 nm, with DiY peak signal expected between 400-420 nm. Tyrosine fluorescence signal was monitored using an excitation wavelength of 280 nm and emission between 290 – 600, with the peak tyrosine emission observed at 305 nm. For experiments involving MCO, the reaction was quenched using EDTA at a final concentration of 2 mM. For all the measurements, the excitation and emission slits were both set to 10 nm with a scan rate set to 300 nm/min with 2.5 nm data intervals and an averaging time of 0.5 s. The photomultiplier tube detector voltage was set at 500 V.

### Thioflavin S (ThS) fluorescence assay to monitor dGAE self-assembly

dGAE samples (10 μL) were mixed and incubated for 3 min with 5 μM ThS in 3-(N-morpholino) propanesulfonic acid (MOPs) (20 mM, pH 6.8 at 37°C). Fluorescence intensity was observed using SpectraMax i3 plate reader. The readings were collected in a black, 96-well plate with a clear bottom (PerkinElmer), which was sealed with an optically clear polyolefin film to avoid evaporation (StarSeal Advanced Polyolefin Film, Starlab). The number of readings per well was set to 6, PMT voltage was set to high and blank spectra of the buffer blank were subtracted from protein fluorescence scans. The excitation wavelength was set at 440 nm, with emission between 460 to 600 nm, with the peak emission observed at 483 nm. Each experiment included a minimum of three independent experiments.

### Circular Dichroism (CD)

The secondary structure of the samples was assessed using Jasco J715 CD spectrometer (Jasco, Goh-Umstadt, Germany). Each sample (40 μL) was placed into a 0.2-mm path length quartz cuvette (Hellma) and scanned between 190 nm and 260 nm. The CD spectra were collected in triplicate at a temperature of 21 °C. Samples were centrifuged at 20,000*g* for 30 min to pellet any fibrillar structures. The pellet was then resuspended in phosphate buffer (pH7.4) before CD analysis. CD data were converted into molar ellipticity (deg·cm^2^·dmol^−1^) where concentrations were known.

### Negative-stain transmission electron microscopy (TEM)

Samples (4 μL) for TEM were placed on 400-mesh carbon-coated grids (Agar Scientific, Essex, UK). After 1 min incubation, excess sample was blotted using filter paper, and the grid was washed with 4 μL filtered Milli-Q water and blotted. The grid was then negatively stained for 40 sec using 4 μL of filtered 2% (w/v) uranyl acetate. The excess stain was blotted with filter paper and grids left to air-dry before storage. The grids were examined on a JEOL Jem1400-plus transmission electron microscope (Jeol, USA), operated at 80 kV fitted with a Gatan Orius SC100 camera (UK).

### Dot immunoblotting

A total of 5◻μl of each sample was spotted onto a 0.2-μM pore nitrocellulose membrane and allowed to dry for 10 min. The membrane was blocked with blocking buffer (5% milk in TBS containing 0.05% Tween 20) for 1◻hour at room temperature on a rocker. The blocking buffer was next replaced with rabbit polyclonal T22 antibody that recognises oligomeric Tau (diluted 1/2000) ((ABN454; Merck Millipore) (37)) and incubated overnight at 4°C on a rocker. The membrane was washed six times for 5 min with washing buffer (0.05% TBS-T), then incubated with an HRP-conjugated goat anti-rabbit secondary antibody for 1 hour. The membrane was washed six times for 5 min with washing buffer, then incubated with Clarity Western ECL Substrate (Bio-Rad) for 1 min before being developed in the darkroom. A minimum of three independent experiments were conducted to ensure reproducibility of the findings.

### TEM Immunogold labelling

DiY was detected using immunogold electron microscopy in the oxidised dGAE sample using methods previously described (23). Briefly, a phosphate-buffered saline, pH 8.2, containing 1% BSA, 500 μl/l Tween-20, 10 mM Na EDTA, and 0.2 g/l NaN3 (henceforth called PBS+), was used throughout. 4 μL of the control and oxidised dGAE samples were placed onto 400-mesh carbon-coated grids (Agar Scientific, Essex, UK), allowed to adhere for 1 min, and the excess sample removed using filter paper. The grids were blocked using normal goat serum (1:10 in PBS+) for 15 min, then incubated with (10 μg/ml IgG) mouse dityrosine monoclonal antibody (JaICA, Shizuoka, Japan) for 2 h at room temperature. The grids were rinsed three times for 2 min in PBS+, and then labelled with a 10 nm gold particle-conjugated goat anti-mouse IgG secondary probe (GaM10 British BioCell International, Cardiff, UK; 1:10 dilution) for 1 h at room temperature. The grids were rinsed 5 times for 2 min using PBS+, 5 times for 2 min with distilled water, then negatively stained as described above.

### Cell death assay

dGAE samples, with or without H_2_O_2_ (Table 1), were exchanged into phosphate buffer (pH 7.4) using disposable Vivaspin® 500 centrifugal concentrators with an MWCO of 3 kDa (Sartorius) to remove Cu^2+^ or H_2_0_2_. Differentiated SHSY5Y neuroblastoma cells were used for the toxicity experiments. Firstly, undifferentiated SHSY5Y neuroblastoma cells (ATCC CRL-2266™), were maintained in Dulbecco’s Modified Eagle Medium: Nutrient Mixture F-12 (DMEM/F-12) (Life Technologies, United Kingdom), supplemented with 1% (v/v) L-glutamate (L-Glu) (Invitrogen), 1% (v/v) penicillin/streptomycin (Pen/Strep) (Invitrogen) and 10% (v/v) Fetal Calf Serum at 37°C and 5% CO_2_. The undifferentiated SHSY5Y cells were seeded to 60% confluency in a CellCarrier-96 Ultra Microplates (PerkinElmer). The cells were differentiated in a medium containing 1% Fetal Calf Serum supplemented with 10 μM trans-retinoic acid (Abcam) for 5 days. Next the medium was replaced with a serum-free media supplemented with 2 nM brain-derived neurotrophic factor (BDNF) (Merck Millipore). After 2 days in the BDNF-containing media, the media was replaced with serum-free media and the cells were treated with phosphate buffer (pH 7.4) or 10 μM of the selected dGAE reaction mixtures for 2 days. At the end of the incubation period, the cells were incubated with ReadyProbes reagent (Life Technologies) for 15 min. The ReadyProbes kit contains NucBlue Live reagent that stains the nuclei of all live cells and Propidium iodide that stains the nuclei of dead cells with compromised plasma membrane. The cells were imaged at 37°C and 5% CO_2_ using Operetta CLS high-content analysis system (PerkinElmer) using DAPI and TRITC filters. At least 5,000 dead and live cells were analysed using the Harmony software automated analysis algorithm within the Operetta CLS high-content analysis system. A minimum of three independent experiments were performed.

## Results

### *In vitro* copper-catalysed oxidation results in the formation of stable, random coil and ThS-negative, dityrosine cross-linked tau oligomers

Different oxidation conditions catalysed by metals (e.g. Cu^2+^), enzymes (e.g. horse-radish peroxidase) and light (e.g. UV) have been used to generate DiY cross-linking in proteins (41). Initially, dGAE was incubated with combinations of Cu^2+^, H_2_O_2_ or EDTA at 37°C as shown in Table 1. DiY is observed as a fluorescent signal with an emission peak between 400–420 nm (23). Fluorescence spectroscopy showed that dGAE incubated with Cu^2+^ at a ratio of 1:10 with 2.5 mM H_2_O_2_ [1-10/Cu^2+^ H_2_O_2_] showed the highest signal at 405 nm following 15 mins of incubation, while Cu^2+^ alone at 1:10 [1-10/Cu^2+^] also showed a lower, but significant signal (Figure 1A). The control sample (buffer only), dGAE incubated with Cu^2+^ at a ratio of 1:0.1 alone [1-0.1/Cu^2+^] or additional 2.5 mM H_2_O_2_ (1-0.1/ Cu^2+^ H_2_O_2_) and dGAE with EDTA alone at 1:1000 ratio (1-1000 EDTA) showed no signal intensity at 405 nm (Figure 1A). This results show that incubation with Cu^2+^ results in the rapid formation of DiY, that is greater in the presence of H_2_O_2_. The presence of DiY was confirmed in the [1-10/Cu^2+^ H_2_O_2_] sample using TEM immunogold labelling using anti-DiY antibody (Fig. 1B). The increase in DiY was matched by a decrease in tyrosine signal at 305 nm for [1-10/Cu^2+^ H_2_O_2_] and [1-10/Cu^2+^] and a strong signal arising from DiY could also be observed in the [1-10/Cu^2+^ H_2_O_2_] sample at 400 nm (Fig. 1C). The [1-0.1/Cu^2+^] and [1-0.1/Cu^2+^ H_2_O_2_] samples that showed no signal for DiY after 15 min incubation, but showed a slight decrease in the tyrosine fluorescence intensity consistent with a conformational change in dGAE (Fig. 1C). After 24h, we observed that the DiY signal intensity reaches a plateau after about 4h incubation in both the [1-10/Cu^2+^] and [1-10/Cu^2+^ H_2_0_2_] samples (Fig. 1D). Together, these data reveal that tau, similar to Aβ (23) and α-synuclein (24), rapidly forms DiY in an MCO environment and that this depends on Cu^2+^concentration. Even in the absence of H_2_O_2,_ supra-equimolar level Cu^2+^ was able to efficiently induce DiY in dGAE.

**Figure 1.**
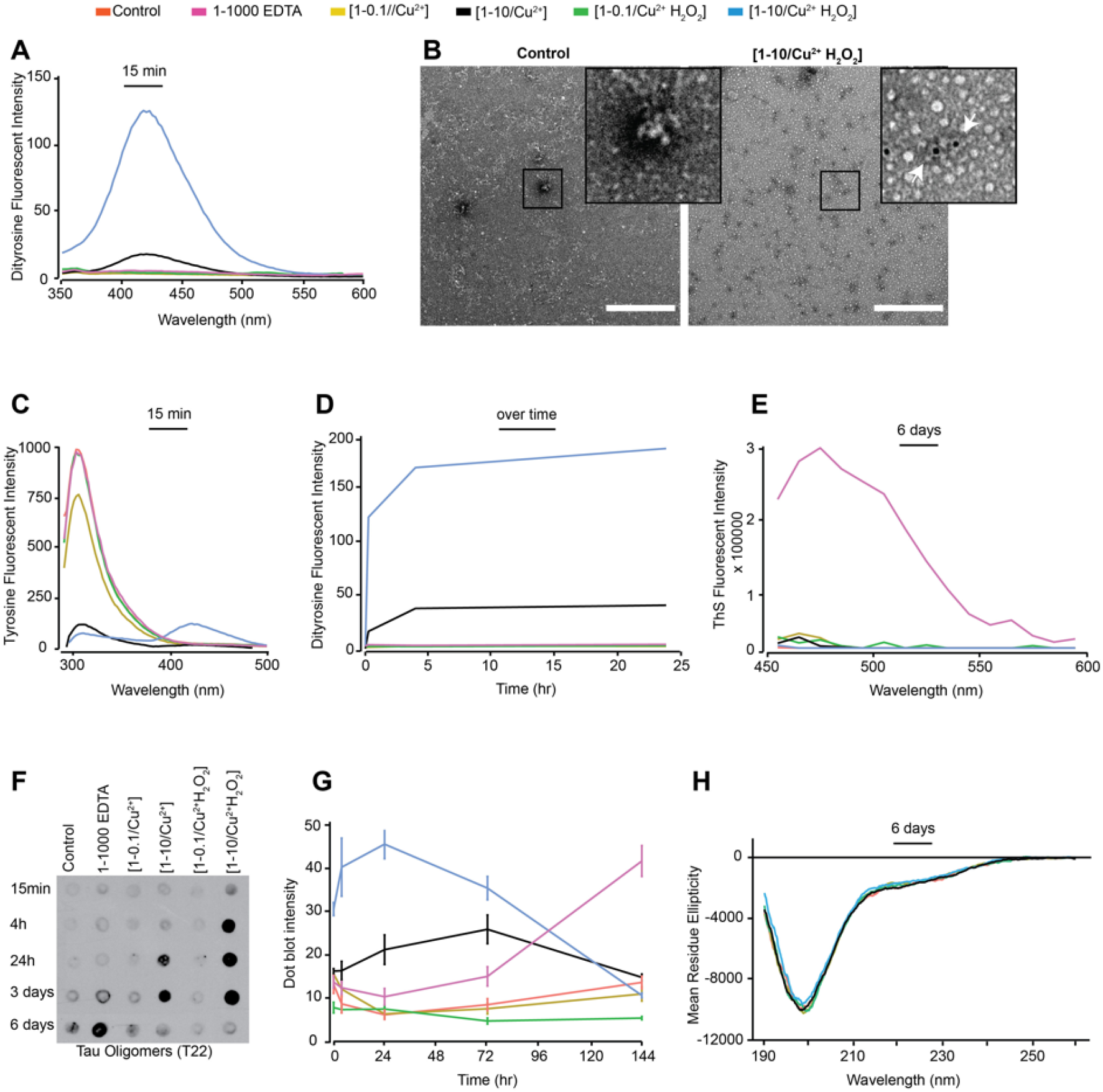
Freshly prepared dGAE (100 μM) was incubated at 37°C without agitation, alone (control), with 10 μM CuCl_2_ [1-0.1/Cu^2+^], 1 mM CuCl_2_ [1-10/Cu^2+^], 10 μM CuCl_2_ in combination with 2.5 mM H_2_O_2_ [1-0.1//Cu^2+^ H_2_O_2_], 1 mM CuCl_2_ in combination with 2.5 mM H_2_O_2_ [1-10/Cu^2+^ H_2_O_2_) or with 100 mM EDTA (1-1000 EDTA). DiY signal was collected 15 min post-incubation using fluorescent excitation/emission 320nm/340 – 600 nm, with DiY peak signal observed between 400-420 nm **(A)**. The presence of DiY in dGAE was confirmed using TEM immunogold-labelling where black dots indicate DiY probed with mouse anti-DiY antibody (see white arrows in the insert). Scale bar equals 500 nm. **(B)**. Tyrosine signal was collected at 15 min of incubation using an excitation/emission 280nm/ 290 – 500nm, with peak tyrosine signal observed at 305 nm **(C).** DiY signal was collected again at 4h and 24h to follow DiY formation over 24h duration (**D)**. ThS fluorescence intensity was conducted after six days incubation to observe the degree of self-assembly using excitation/emission of 440/460-600nm (**E**). Immunoblotting using T22 antibody identifies the presence of tau oligomers (**F**). Quantification of the immunoblot signal suggest a high level of tau oligomers early-on in the [1-0.1//Cu^2+^ H_2_O_2_]and [1-0.1//Cu^2+^] samples (**G**). CD analysis at six days post-incubation showed that all the samples had a comparable level of random-coil secondary structure (**H**).

Thioflavin S (ThS) fluorescence assay was used to investigate the assembly of dGAE under the various conditions. ThS fluorescence intensity increased only in the 1-1000 EDTA sample suggesting that dGAE assembles only in the presence of the metal chelator, while all other conditions showed very low fluorescence of ThS at 483 nm up to six days suggesting that dGAE does not assemble in the absence of agitation (Fig. 1E). dGAE samples incubated with lower concentrations of EDTA (1-10 and 1-100) did not show ThS fluorescence (not shown).

Immunoblotting was conducted using T22, the tau oligomer-specific polyclonal antibody, to learn more about the conformation of dGAE in the different conditions (42). Interestingly, immunoblots showed higher intensity suggesting a higher level of tau oligomers from 4h to 3 days in the samples that also developed DiY, compared to those samples in which DiY did not form (Fig. 1F-G). [1-10/Cu^2+^ H_2_O_2_], which favours rapid DiY formation, showed strong T22 binding affinity as early as 15 min post-incubation which remained intense up to 3 days. [1-10/Cu^2+^] sample showed significant antibody binding intensity at 24h, which further increased at 3 days following incubation (Fig. 1F-G). The T22 binding affinity for both the [1-10/Cu^2+^] and [1-10/Cu^2+^ H_2_O_2_] samples had decreased after six days suggesting that the DiY formation facilitates the formation of T22-positive but ThS negative tau oligomers and prolonged their half-life. The disappearance of the T22 signal after this time maybe due to further oxidation of the dGAE.

We have previously shown that soluble dGAE exists in a random-coil conformation by circular dichroism (CD), and transitions to β-sheet conformation during self-assembly, accompanied by a reduction in random coil signal (33). Moreover, tau oligomers have been described as being insoluble with β-sheet secondary structure (37). We used CD to interrogate the secondary structure of the dGAE under the different conditions, and this revealed that all the samples remained in a random-coil conformation with an identical CD spectrum (Fig. 1H). Given that tau exists mostly in a random-coil conformation, CD spectra analysis can be conducted by resuspending a pellet obtained from high-speed centrifugation of the dGAE samples in order to investigate the presence of a β-sheet signal (33). However, none of the samples, except the 1-1000 EDTA reaction, formed a significant pellet after centrifugation, which precluded the search for β-sheet signal in these samples. The high concentration of EDTA in the 1-1000 EDTA reaction led to a significant increase in the high-tension signal during CD spectra collection (not shown). Therefore, CD could not be conducted on the 1-1000 EDTA sample. Together, the CD data suggests that oxidation causes no obvious change in secondary structure or solubility of the dGAE under any of the conditions except in high EDTA.

TEM negative staining was used to examine the morphology of dGAE in the different samples (Fig. 2). At 0h, dGAE under all conditions showed the typical morphology of unassembled dGAE (33). By 6 days, however, the control sample revealed the presence of small assemblies, which are mostly short rod-like particles, likely on the path to becoming mature fibrils. The 1-1000 EDTA sample showed a combination of short and long fibrils, consistent with the ThS data indicating self-assembly. [1-0.1/Cu^2+^] and [1-0.1/Cu^2+^ H_2_O_2_] samples both showed small assemblies and short fibrils. Interestingly, the [1-10/Cu^2+^] and [1-10/Cu^2+^ H_2_O_2_] samples, which both form DiY, showed very few short fibres and large amorphous aggregates. In summary, TEM data revealed that only the 1-1000 EDTA reaction goes on to form long tau filaments after six days incubation, while the other samples form assemblies that are largely non-fibrillar and consistent with the observation of very low ThS fluorescence. The data suggest that DiY cross-linking results in oligomerisation and stabilisation of dGAE into large oligomers that are rich in random-coil.

**Figure 2.**
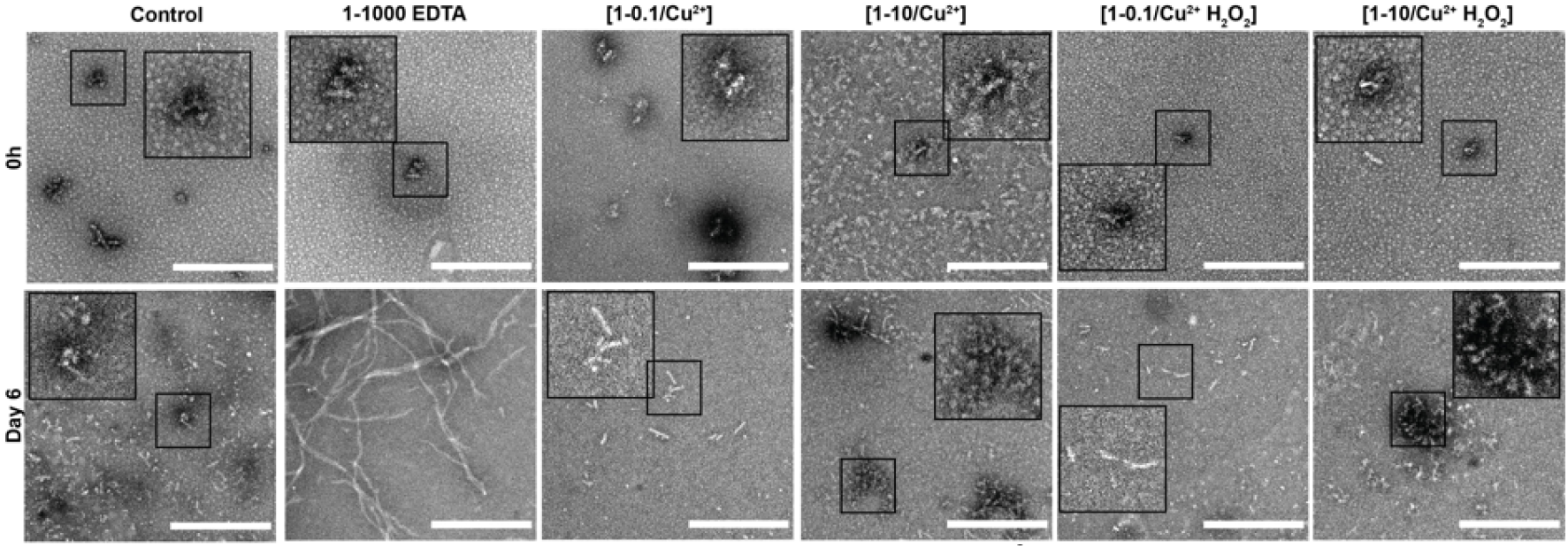
Freshly prepared dGAE (100 μM) was incubated at 37°C without agitation, alone (control), with 10 μM CuCl_2_ [1-0.1/Cu^2+^], 1 mM CuCl_2_ [1-10/ Cu^2+^], 10 μM CuCl_2_ in combination with 2.5 mM H_2_0_2_ [1-0.1/Cu^2+^ H_2_O_2_], 1 mM CuCl_2_ in combination with 2.5 mM H_2_0_2_ [1-10/ Cu^2+^ H_2_O_2_] or with 100 mM EDTA (1-1000 EDTA). TEM imaging of all the samples at 0h revealed small, round assemblies (**Top row**). At six days, the control, [1-0.1/Cu^2+^] and [1-0.1/Cu^2+^ H_2_O_2_] samples show the presence of small, short fibrillar assemblies. The [1-10/Cu^2+^] and [1-10//Cu^2+^ H_2_O_2_] samples showed small and large oligomeric assemblies with the occasional presence of short fibrils. The 1-1000 EDTA samples revealed short and long fibrils. Scale bar set at 500 nm.

### Oxidation resulting in DiY cross-linking halts elongation of tau oligomers to fibrils following agitation

We have previously shown that the agitation of dGAE enhances its self-assembly into filaments (33) shortening the incubation time to 24h. Therefore, 100 μM dGAE was prepared as described earlier (Table 1), except that the samples were incubated at 37°C in a thermomixer set to oscillate at 350 RPM to facilitate assembly. The 37°C/350RPM condition results in DiY formation in both the [1-10/Cu^2+^] and [1-10/Cu^2+^ H_2_O_2_] samples over time (Fig. 3A, C). However, the DiY signal was slightly higher than the signal observed in similar reaction mixtures incubated without agitation (Fig. 1A, D). As with the result obtained with the quiescent condition (Fig. 1C), agitation led to a decrease in the tyrosine signal in the [1-10/Cu^2+^] and [1-10/Cu^2+^ H_2_O_2_] samples with incubation time (Fig. 3B). Agitation has been shown to enhance assembly and, as expected, all conditions except the [1-10/Cu^2+^] and [1-10/Cu^2+^ H_2_O_2_] led to ThS intensity increase over time (Fig. 3D). [1-10/Cu^2+^] and [1-10/Cu^2+^ H_2_O_2_] conditions did not show any increase in ThS fluorescence even after 150 hours. These data support the view that oxidation, which results in formation of DiY, inhibits further elongation (Fig. 1E), and suggests that those reactions in which DiY forms ([1-10/Cu^2+^] and [1-10//Cu^2+^ H_2_O_2_]) have reduced self-assembly properties.

**Figure 3.**
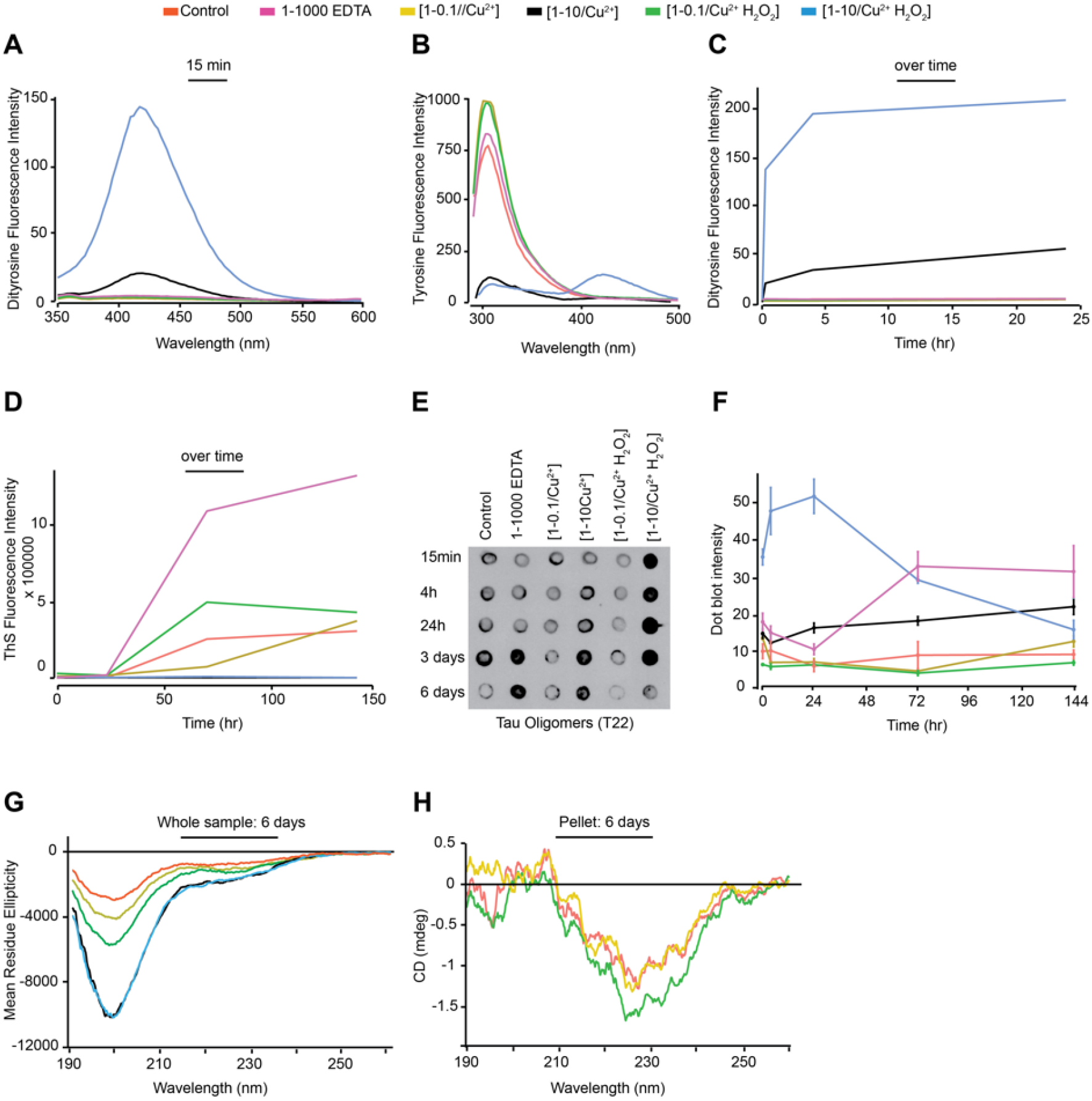
Freshly prepared control, [1-0.1/Cu^2+^], [1-10 Cu^2+^], [1-0.1/Cu^2+^ H_2_O_2_], [1-10/Cu^2+^ H_2_O_2_] and 1-1000 EDTA samples were incubated at 37°C/350 RPM. DiY fluorescence signal was collected 15 min post-incubation, showing a significant increase in intensity at 410-420 nm in the [1-10/Cu^2+^] and [1-10/Cu^2+^ H_2_O_2_] samples **(A)**. Tyrosine signal was collected at 15 min of incubation showing a significant decrease in intensity over time for the 1-10 and 1-10/H_2_O_2_ samples **(B)**. DiY signal was collected at 4h and 24h to follow DiY formation over 24h duration, which showed that DiY reaches plateau about 4h after incubation in the [1-10/Cu^2+^ H_2_O_2_], while it continues to increase in the [1-10/Cu^2+^] **(C)**. ThS fluorescence was collected over a time course of six days to assess the degree of self-assembly over time (**D**). Immunoblotting using T22 antibody reveals the presence of Tau oligomers (**E**). Quantification of the dot blotting signal revealed a high level of Tau oligomers early-on in the [1:10/Cu^2+^ H_2_O_2_] reaction, followed by a gradual increase in the oligomer levels in the [1:10/Cu^2+^] reaction (**F**). CD at six days showed a reduction in the level of random-coil in control, [1:0.1/Cu^2+^], [1:0.1/Cu^2+^ H_2_O_2_] samples when compared to the [1:10/Cu^2+^] and [1:10/Cu^2+^ H_2_O_2_] samples (**G)**. The CD spectra for the pellet from the control, [1:0.1/Cu^2+^], [1:0.1/Cu^2+^ H_2_O_2_] samples revealed the presence of a minimum at 228nm (**H**).

T22 immunoblotting showed that [1-10/Cu^2+^] and [1-10/Cu^2+^ H_2_O_2_] at 37°C/350RPM showed increased T22 binding intensity at early time points which remained high up to three days incubation (Fig. 3E-F) The levels of T22 binding were especially very high in the [1-10/Cu^2+^ H_2_O_2_] 37°C/350RPM sample (Fig. 3F). These data support the view that DiY cross-linking promotes the formation of long-lived, soluble tau oligomers that show negilible ThS fluorescence.

CD showed the expected reduction in random coil signal at 195nm over time as the dGAE assembles under all conditions except for samples containing [1-10/Cu^2+^] and [1-10/Cu^2+^ H_2_O_2_] (Fig. 3G). Samples were centrifuged to separate the pellet and supernatant, and the control, [1-0.1/Cu^2+^] and [1-0.1/Cu^2+^ H_2_O_2_] samples showed a small minimum at 228 nm consistent with previous observations attributed to elongated β-sheet (33) (Fig. 3H). However, even after the 30 min high-speed centrifugation, the [1-10 Cu^2+^] and [1-10/ Cu^2+^ H_2_O_2_] samples did not form pellets. CD could not be obtained from the 1-1000 EDTA sample due to the EDTA concentration in the sample that caused an increase in the high-tension signal, which makes the result unreliable. CD spectra analysis agrees with the ThS assay results in showing that the ThS-positive dGAE reactions are β-sheet positive, while the ThS-negative samples are random-coil rich.

TEM negative staining at 0h showed the typical morphology of unassembled dGAE for all samples (Fig. 4). However, following six days incubation at 37°C/350RPM, all the samples except the 1-10 and 1-10/H_2_0_2_ samples, formed filaments consistent with the results from ThS fluorescence (Fig. 3D). The 1-1000 EDTA sample showed the highest density of fibres, comprised of both short and long, twisted fibres. In contrast, oxidised DiY cross-linked [1-10/Cu^2+^] and [1-10/Cu^2+^ H_2_O_2_] dGAE showed mainly amorphous oligomeric aggregates (Fig. 2). Altogether, this data suggests that MCO-induced DiY cross-linking promotes the assembly and stabilisation of the dGAE into ThS-negative, random-coil rich, oligomeric assemblies and prevents further elongation into filaments.

**Figure 4.**
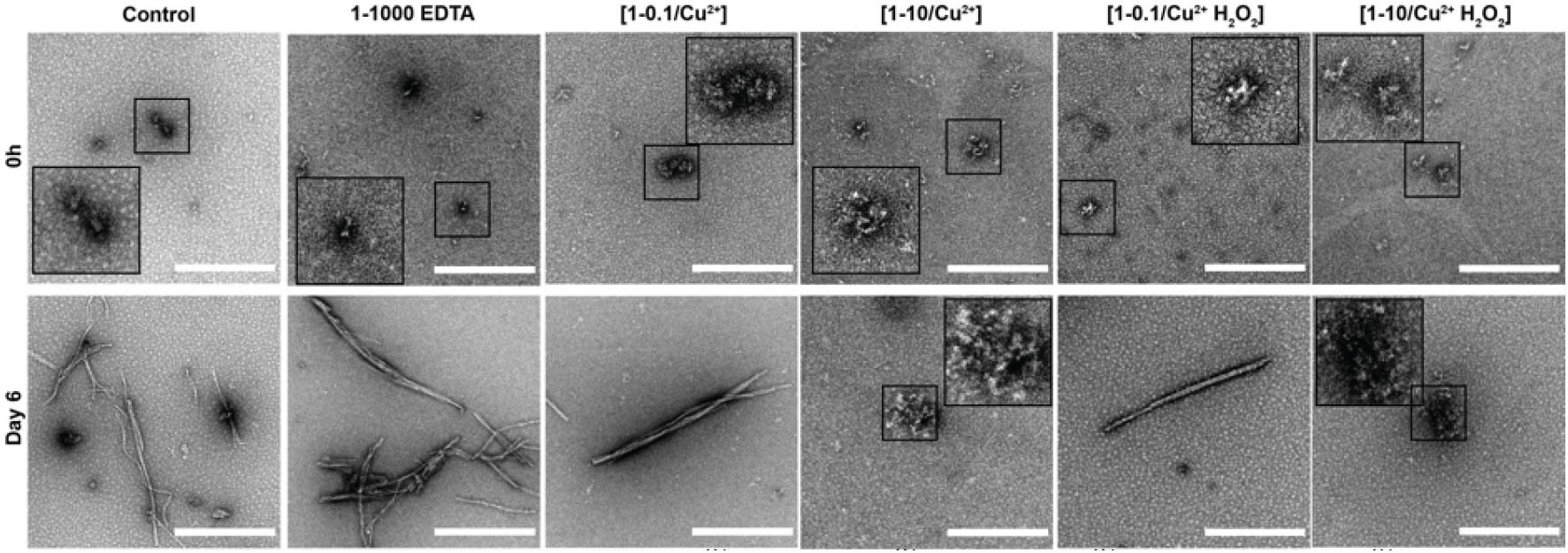
Freshly prepared control, [1-0.1/Cu^2+^], [1-10/Cu^2+^], [1-0.1/Cu^2+^ H_2_O_2_], [1-10/Cu^2+^ H_2_O_2_] and 1-1000 EDTA samples were incubated at 37°C/350 RPM. TEM imaging of all the samples at 0h revealed small, round assemblies (**Top row**). At six days post-incubation, the control, [1-0.1/Cu^2+^] and [1-0.1/Cu^2+^ H_2_0_2_] samples showed both short and long fibrillar assemblies. The [1-10/Cu^2+^] and [1-10/Cu^2+^ H_2_0_2_] samples showed mostly small and large amorphous assemblies. The 1-1000 EDTA samples revealed mature, twisted filaments. Scale bar equals 500 nm.

### DiY formation before the onset of assembly promotes the formation of tau oligomers but inhibits elongation into fibrils

We have shown that DiY cross-linking, induced by supra-equimolar Cu^2+^ and CuCl_2_ with H_2_0_2_, prevents elongation of dGAE into filaments. Previous studies have shown that Cu^2+^ influences tau self-assembly (43, 44). Therefore, to evaluate the impact of DiY on dGAE assembly without the influence of agitation, temperature or Cu^2+^, UV photo-oxidation was performed at a low temperature and without agitation, to induce DiY cross-linking. We first optimised the UV exposure duration to induce DiY formation in dGAE. UV exposure for 2h at 4°C resulted in the formation of DiY in a freshly prepared dGAE sample (+2h UV) (Fig. 5A). DiY intensity was ~ 5-fold lower than the DiY signal observed after 15 min in the [1-10/Cu^2+^ H_2_O_2_] incubated at 37°C/350 RPM (Fig. 3A). When the same dGAE sample incubated with UV for 2h was further incubated at 37°C/350 RPM for six days, the level of DiY doubled (+2h UV/6d) (Fig. 5A). ThS fluorescence assay indicated that the UV exposure did not result in dGAE self-assembly after six days incubation following 2h UV exposure similar to the control (Fig. 5B). Immunoblotting using the T22 antibody showed that the 2h UV-exposure results in the formation of T22-positive oligomers which remains after six days incubation (Fig. 5C). Therefore, 2h UV exposure resulted in formation of DiY, inhibition of assembly and stabilisation of T22 positive oligomers which were maintained for six days after UV incubation.

To investigate whether addition of EDTA can modulate the DiY formation and assembly of UV incubated dGAE, a freshly prepared 1-1000 EDTA sample was incubated for 2h under UV [1-1000 EDTA +UV] and compared to EDTA without UV [1-1000 EDTA] (Fig. 5D). A DiY signal was observed after 2h UV incubation of dGAE in the presence of EDTA and increased after 6 days agitation at 37°C while those unexposed to UV did not form DiY (Fig. 5D). As expected, the sample that was not exposed to UV showed significant ThS fluorescence after 6 days incubation at 37°C/350 RPM and only a very small signal was observed for the 6day EDTA dGAE sample incubated in UV (Fig. 5E). Electron micrographs showed that the expected filaments were present in dGAE incubated with EDTA (Fig. 5F) but the UV-exposed EDTA sample contained only amorphous assemblies even after six days of incubation at 37°C/350 RPM (Fig. 5G). DiY formation before the onset of assembly inhibits or substantially delays the assembly of the dGAE into filaments.

### DiY cross-linked tau oligomers are not toxic to cells

The evidence for neurotoxic assemblies formed from tau remains unclear (5). Toxic tau oligomers have been described as having β-sheet secondary structure (37) while non-toxic, β-sheet negative tau oligomers have also been described (45). To investigate and compare the toxicity of non-oxidised and DiY dGAE, we utilised differentiated human neuroblastoma SHSY5Y cells (15, 46) and assessed the effect on cell survival using the ReadyProbes cell viability assay. We have established that Cu^2+^ and Cu^2+^ with H_2_O_2_, as well as 2h UV induce significant DiY cross-linking, resulting in long-lived amorphous oligomers (Fig. 1–5). Differentiated SHSY5Y cells were incubated for 72 hours with phosphate buffer (as vehicle control) or 10 μM 2h UV-exposed dGAE, 10 μM 3 days/37°C-incubated dGAE control, [1-10/Cu^2+^] sample and [1-10/Cu^2+^ H_2_O_2_] sample. No significant cell death was detected for any of the conditions. Incubation for 1h with 2 mM H_2_0_2_ was used as a positive control (Fig. 6). These data revealed that the DiY cross-linked, ThS and β-sheet negative soluble tau oligomers described here are not acutely toxic to cells.

**Figure 5.**
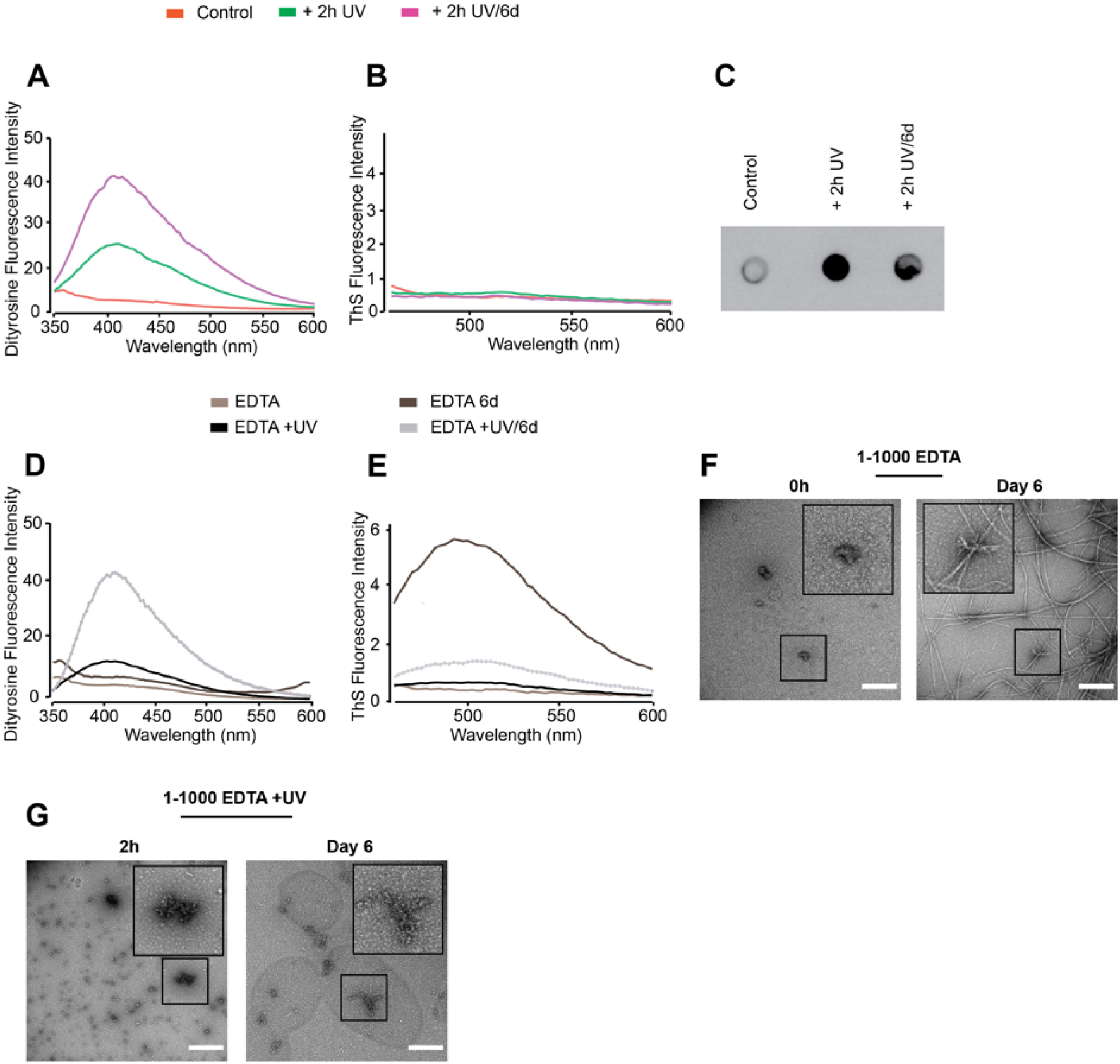
Freshly prepared dGAE samples (100 μM) were incubated without UV (control) or in the presence of UV 2h at 4°C [+2h UV], and the 2h UV-exposed sample was further incubated for 6days at 37°C/350 RPM [+2h UV/6d]. DiY signal over time and revealed the induction of DiY by the UV exposure (**A**). ThS fluorescence assay showed no fluorescence intensity indicating there was no assembly in any of the samples (**B**). Immunoblotting using the T22 antibody, suggested the presence of tau oligomers in a high quantity in the UV-exposed samples compared to control (**C**). Freshly prepared 1-1000 EDTA samples were incubated without UV for 2h or 6days at 37°C/350 RPM (1-1000 EDTA 6d) and with UV for 2h (1-1000 EDTA +UV) or 2h UV followed by 6days incubation at 37°C/350 RPM (1-1000 EDTA +UV/6d). DiY signal was collected at each time points (**D**), followed by ThS fluorescence assay, which suggested assembly in the 1-1000 EDTA 6d sample and not the others (**E**). TEM imaging at 0h revealed small, round assemblies in the 0h [1-1000 EDTA] sample, which assembled into long mature fibrils after 6days incubation at 37°C/350 RPM (**F**). The 1-1000 EDTA + UV samples revealed small and large clumped assemblies, which remained largely unchanged even after 6days of incubation at 37°C/350 RPM. Scale bar set at 500 nm.

**Figure 6.**
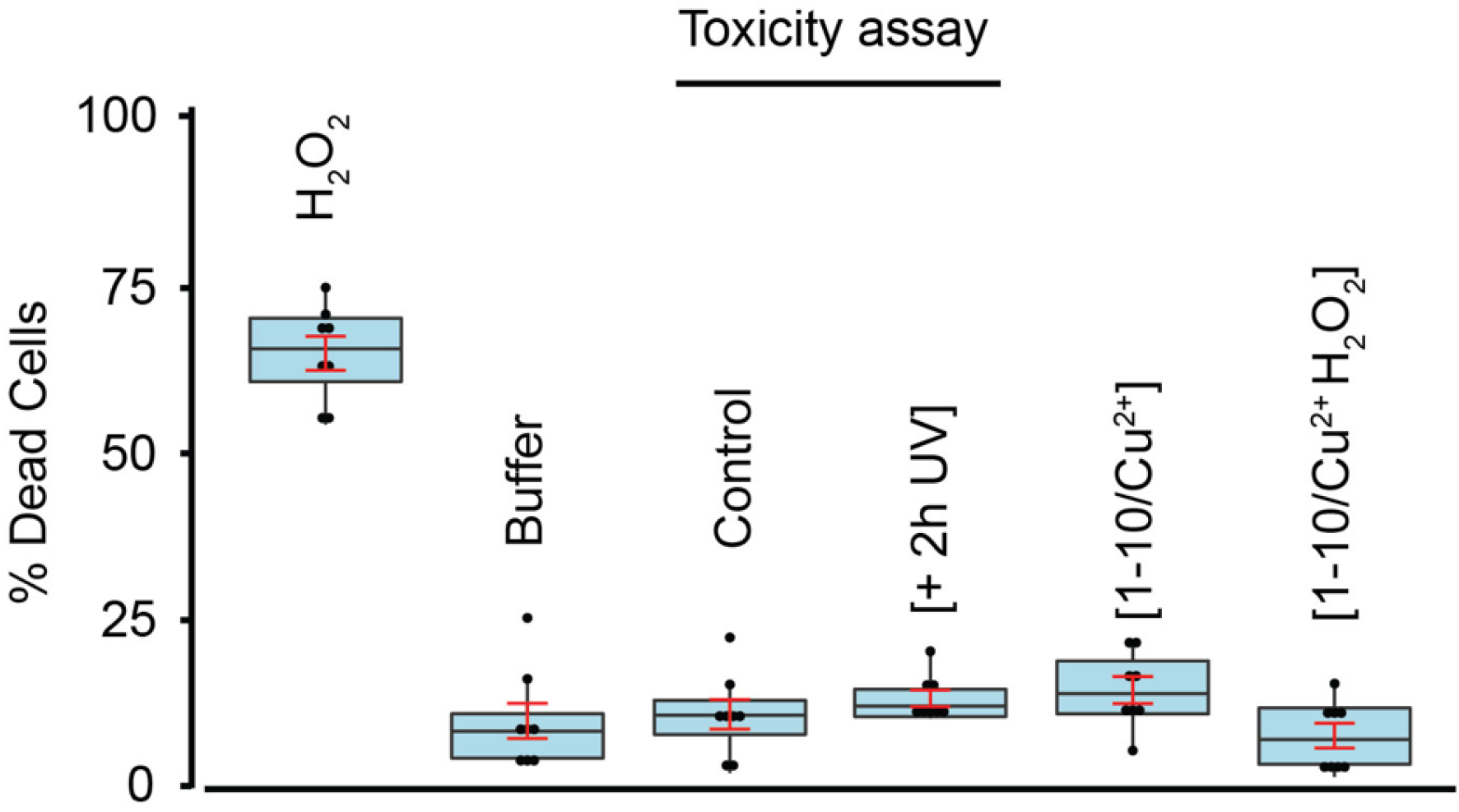
Cellular toxicity. Differentiated SHSY5Y cells were incubated for three days in the presence of phosphate buffer, or 10 μM 2h UV-exposed dGAE, 10 μM three days/37°C-incubated dGAE control, [1-10/Cu^2+^] sample and [1-10/Cu^2+^ H_2_0_2_] sample. 1h incubation of cells with 2 mM H_2_0_2_ was used as a positive control. At the end of the incubation period, cells were incubated with ReadyProbes reagent for 15 min, imaged at 37°C and 5% CO_2_ using Operetta CLS high-content analysis system, and at least 5,000 dead and live cells were used for analysis. Only cells treated with 2 mM H_2_0_2_ showed a significant % of dead cells compared to control.

## Discussion

Oxidative stress has been proposed to play a key role in AD (47, 48) and it has been suggested that it may be among the earliest sources of damage in human AD (18). The levels of nitrotyrosine and DiY – two products of oxidative modification, have been shown to be significantly increased in the AD brain (19). The cross-linking of proteins by DiY enhances their stability (20) demonstrated by the occurrence in tough proteins such as resilin. Indeed, DiY cross-links occur naturally in several elastic and structural proteins including elastin, fibroin, keratin, cuticlin, and collagen (49–52). In these proteins, DiY cross-links can contribute to increased mechanical strength and protein insolubility (53). DiY cross-linking has been shown to form and influence the aggregation of both Aβ and α-synuclein *in vitro* under oxidative environments (21–23, 54). We have previously demonstrated that DiY cross-linking stabilises Aβ and α-synuclein fibrils *in vitro* and revealed the presence of DiY-cross-linked Aβ fibrils in Aβ plaques and DiY cross-linked α-synuclein in Lewy bodies (23, 24). These findings suggested that oxidative stress influences Aβ and α-synuclein assembly and stability via DiY cross-linking but the role of DiY on tau assembly remains unclear. Here, we have explored this question and revealed that the region of tau that forms the core of PHFs, dGAE, is able to form DiY via Y310 *in vitro*. We show that DiY formation facilitates the formation of non-toxic, soluble tau oligomers and inhibits the elongation of the oligomers to fibrils. DiY was induced rapidly in dGAE either via Cu^2+^-catalysed oxidation or UV photo-oxidation. Interestingly, DiY cross-linking facilitated the formation of T22-positive tau oligomers in a DiY-dependent manner. CD, ThS fluorescence and TEM revealed that the DiY cross-linked dGAE oligomers are random-coil rich, ThS negative, amorphous aggregates and are unable to elongate to form fibrils. Previous studies revealed that oxidative stress induced by ONOO^−^ results in the oligomerisation of full-length human tau stabilised via DiY cross-linking (12).

Multiple strands of evidence support the idea that metal ions may play a role in AD pathogenesis. The aberrant distribution of copper, iron and zinc has been shown in the AD brain (55). For example, ~ 400 μM copper has been estimated around Aβ plaques (55). There is substantial evidence that copper can influence Aβ and tau aggregation (43, 44) and efficiently catalyse DiY cross-linking on Aβ and α-synuclein (23, 24). Copper can bind tau via its microtubule-binding repeat region (43, 56), which is contained in the dGAE fragment used in this study (33). Here, we showed that at a supra-equimolar ratio, Cu^2+^ facilitates DiY cross-linking and tau oligomerisation, prolongs the oligomer half-life and inhibits the further assembly of the oligomers into fibrils. While at a sub-equimolar ratio, it appears not to influence dGAE assembly. Indeed, multiple studies have studied the influence of copper on tau aggregation and this is still not fully understood. The impact of copper on tau appears to depend on the tau isoform, pH, temperature and other factors (43, 57–59). It would be interesting for future studies to characterise the concentration-dependent role of Cu^2+^ on tau more fully in *in vitro* and *in vivo* environments.

DiY cross-linking has been demonstrated in preformed tau fibrils, leading to the suggestion that the cross-linking may be essential for tau filament assembly (60). Here we have shown that that DiY cross-linking before the onset of assembly promotes tau oligomerisation, but inhibits further elongation. By TEM, the DiY cross-linked dGAE assemblies showed large clumped aggregates that appear to be composed of amorphous oligomers and this is supported by the observation of random coil conformation retained in these samples by CD. This may suggest that the cross-linking traps the oligomers together in a conformation that does not favour further elongation to fibrils. Similarly, cross-linking in tau filaments may stabilise the assemblies (60). However, the nature of tau fibrils and accessibility of the tyrosine residues will be critical for ability to form DiY cross-links. In the core of PHFs and straight filaments, Y310 is buried in one of the eight β-sheets that run along the length of the protofilament, adopting a C-shaped architecture (31). If the tyrosine residues are not exposed, DiY formation may be impeded. Nonetheless, given that there are several tyrosine residues located at residues 18, 29, 197, 310, and 394, DiY could still form with the other tyrosine residues, which would lead to enhanced stability and increased insolubility of PHFs (60). PHFs derived from the AD brain demonstrate striking insolubility and resistance to proteolytic cleavage. Early-stage PHF-tau has reduced SDS solubility (61, 62), while late-stage PHF tau exhibits SDS and sarcosyl insolubility (62–64). Thus, it has been proposed that DiY cross-linking is important for the stabilisation of the early-stage PHF-tau, which confers stability to the PHFs and its conversion into late-stage PHF (60). Overall, our results suggest that DiY can influence the assembly process of tau, but that oxidation at the early stages of assemble result in formation of oligomers that do not progress to form PHFs.

Multiple studies have reported the toxic properties of tau oligomers (37–40). Here, tau oligomers formed as a result of the DiY cross-linking are ThS-negative and random-coil rich, unlike typically studied tau oligomers which are thought to be insoluble and β-sheet rich (37, 65). The DiY cross-linked dGAE oligomers were non-toxic to differentiated neuroblastoma cells during the incubation period studied. This suggests that these oligomers either do not cause cell death under these conditions, or that they cause more subtle forms of neuronal dysfunction not measured here. For instance, it was shown that low-n tau^RDΔK^ oligomers selectively impaired spine morphology and density, accompanied by increased reactive oxygen species and intracellular calcium, but without affecting cell viability (66). However these oligomers were ThS-positive and have minimal β-sheet conformation (66), unlike the DiY cross-linked oligomers reported here which are ThS-negative and random-coil rich. This may indicate that the DiY cross-linking facilitates the formation of non-toxic, off-pathway tau oligomers. Previous work has shown that the rescue of tau toxicity in *Drosophila* results in the formation of non-toxic tau oligomers lacking β-sheet (45). Similarly, it has also been demonstrated that the inhibition of tau aggregation using phthalocyanine tetrasulfonate (PcTS) results in the formation of β-sheet negative, soluble tau oligomers (67). In the latter study, it was shown that PcTS interacts with tyrosine residues on tau, including Y310 to induce its assembly into oligomers. This is interesting given that the DiY cross-linking observed in our work occurs via Y310 since it is the only tyrosine in the dGAE fragment. This suggests that the accessibility of this residue may be involved in the generation of this β-sheet negative oligomeric tau species. Overall, these data further confirm the diversity of tau oligomers and that the structural conformation of the oligomers may be key to their toxicity.

In conclusion, our findings suggest that DiY formation facilitates the oligomerisation of dGAE, but inhibits or significantly delays its elongation into fibrils. We have shown that DiY cross-linked soluble dGAE oligomers do not have features of amyloid aggregates and are not toxic. This finding has implications for understanding the toxic species of tau and therapeutic approaches aimed at inhibiting tau aggregation.

## Acknowledgements

The authors acknowledge the Electron microscopy imaging centre of the University of Sussex, funded by the School of Life Sciences, the Wellcome Trust (095605/Z/11/A, 208348/Z/17/Z) and the RM Phillips Trust for their support & assistance in this work. This work was supported by funding from Alzheimer’s Society [345 (AS-PG-16b-010)] awarded to LCS and funding MBM. YA is supported by WisTa Laboratories Ltd (PAR1596). The work was supported by ARUK South Coast Network. GB was supported by European Molecular Biology Organisation (EMBO) Short-Term Fellowship award (EMBO-STF 7674). LCS is supported by BBSRC [BB/S003657/1].

## Author contributions

MBM planned and carried out the work. YA and GB contributed experimental work. MBM and LCS wrote the paper. YA, CRH and CMW reviewed and edited the paper. LCS managed the project.

MBM is supported by Alzheimer’s Society project grant awarded to LCS. GB is supported by EMBO fellowship. LCS is supported by Alzheimer’s Research UK and Alzheimer’s Society.

